# Untangling an insect virome from its endogenous viral elements

**DOI:** 10.1101/2023.05.23.541933

**Authors:** Paula Rozo-Lopez, William Brewer, Simon Käfer, McKayla M. Martin, Benjamin J. Parker

## Abstract

Insects are an important reservoir of viral biodiversity, but the vast majority of viruses associated with insects have not been discovered. Recent studies have therefore employed high-throughput sequencing of RNA, which has led to rapid advances in our understanding of insect viral diversity. However, insect genomes frequently contain transcribed endogenous viral elements with significant homology to exogenous viruses, complicating the use of RNAseq for viral discovery. In this study, we use a multi-pronged sequencing approach to study the virome of an important agricultural pest and prolific vector of plant pathogens, the potato aphid *Macrosiphum euphorbiae*. We first used rRNA-reduced RNAseq to characterize the bacteria and viruses found in individual insects. We then characterized the frequency of a heritable Flavivirus and an Ambidensovirus in our population. We next generated a quality draft genome assembly for *M. euphorbiae* using Illumina-corrected Nanopore sequencing. This analysis showed that the Ambidensovirus, previously described from an RNAseq viral screen, is not a exogenous virus and instead is a transcribed endogenous viral element in the *M. euphorbiae* genome. Our study generates key insight into an important agricultural pest and highlights a widespread challenge for the study of viral diversity using RNAseq.

## Introduction

The last decade has transformed our understanding of the viral communities associated with insects, the most abundant and diversified animal group [1-4]. Insect viruses have been primarily studied in the context of vector-borne pathogens, which are transmitted horizontally between insect vectors and amplifying hosts and often have medical or agricultural relevance. Other viruses, however, only replicate within the insect and are maintained in natural populations through horizontal and/or vertical transmission. These insect-specific viruses have been shown to have important impacts on host biology [5-7], but much work remains to be done to describe insect-specific viral diversity and uncover the hidden role they play in insect phenotypes and evolution [8-10].

To address this gap, researchers have employed high-throughput approaches to viral discovery, including next-generation sequencing and analysis of RNA. Recent studies have used this approach to characterize and discover an enormous diversity of viruses [2, 11-15]. However, there are several serious limitations to this approach. For example, it is unclear from RNAseq data whether viral reads come from microbes infecting insect cells or if they are present from an organism ingested by the insect. Another potential challenge with using RNAseq for viral discovery is that insects often harbor fragments of viral sequences in their genomes. The endogenous viral elements (EVEs) described to date have homology with multiple clades of single- and double-stranded DNA and RNA viral families [16]. We have a limited understanding of the role EVEs are playing in insect biology, but transcriptionally active EVEs have been shown to play functional roles in regulating host genome stability and as an antiviral defense against exogenous viruses [17-19]. EVEs are remarkably common across insects [20], and thus EVEs could represent a widespread challenge facing the field. As such, studies are needed to uncover the contribution of EVEs to insect ‘viromes’.

Aphids (Hemiptera: Aphidoidea) are hosts to diverse viruses, including plant pathogens with agricultural significance and insect-specific viruses [21, 22]. Recent studies have used metatranscriptome sequencing to describe viral diversity in aphids [23-27], and have described insect-specific DNA viruses in the family Parvoviridae (Ambidensovirus) and RNA viruses in the Bunyaviruses, Dicistroviruses, Flaviviruses, Iflaviruses, and Mesoviruses families [21]. The potato aphid *Macrosiphum euphorbiae* (Thomas, 1878) is an important cosmopolitan agricultural pest that infests tomatoes, potatoes, and other economically important crops [28]. *M. euphorbiae* is also an important vector of plant viruses (Families Bromoviridae, Closteroviridae, Geminiviridae, Potyviridae, and Solemoviridae) and was recently shown to host several insect-specific viruses belonging to the families Flaviviridae (Flavivirus) and Parvoviridae [24, 29, 30]. Despite *M. euphorbiae’s* economic importance, no genomic resources are available outside the body and salivary gland transcriptomes [24, 31, 32].

The genomes of multiple aphid species have been shown to harbor EVEs that mediate growth, development, and wing plasticity [33-37]. In this study, we use next-generation sequencing and molecular techniques to show that aphid EVEs have led to the misidentification of aphid viruses from RNAseq data. First, we used RNAseq to characterize the microbial diversity of field-collected *M. euphorbiae* adults, and we found evidence of two insect-specific viruses infecting aphids collected from the field, including a Flavivirus and Ambidensovirus. Then, we generated a high-quality draft genome sequence of this species. Our genome showed that insect-specific Ambidensoviral hits correspond to transcriptionally active EVEs, indicating that a previously described virus is actually an endogenous viral element in the *M. euphorbiae* genome. Our results illustrate how careful analysis using multiple methods is needed to untangle insect viromes from EVEs, and this study furthers our understanding of the surprisingly widespread presence of densoviral EVEs in aphid genomes.

## Methods

### Aphid collection

We collected asexual winged and wingless female *M. euphorbiae* adults from cultivated tomato plants (var Husky Cherry Red) in Knoxville, TN, USA, between April and June 2021 and 2022. We stored individual aphids in 1.5 mL Eppendorf tubes (Eppendorf, Hamburg, Germany) at -80°C until processing. To validate our ability to identify *M. euphorbiae* (NCBI TaxID: 13131), we used COI barcoding (LCO1490 5’-GGTCAACAAATCATAAAGATATTGG-3’ and HCO2198 5’-TAAACTTCAGGGTGACCAAAAAATCA-3’), sanger sequencing, and comparisons of our COI sequences to the Barcode of Life Data System (https://www.boldsystems.org/) [38]. Our partial COI barcode sequence was uploaded to NCBI with accession number OQ588703.

### Cultivation of *M. euphorbiae* strain Me57

To establish a colony of *M. euphorbiae* in the laboratory, we used a single asexual female collected in 2021. After colonization, we maintained this line on tomato plants (Husky Cherry Red) at 20°C 16L:8D. We screened the line for the seven species of facultative symbionts found in aphids using established PCR protocols [39, 40]. For this screen, we extracted DNA using ‘Bender buffer’ and ethanol precipitation as in previous studies [41, 42]. We then used PCR with species-specific primers [39, 43] to screen for *Hamiltonella defensa, Fukatsuia symbiotica* (X-type), *Regiella insecticola, Rickettsia* sp., *Ricketsiella* sp., *Serratia symbiotica*, and *Spiroplasma* sp. following the recommended thermal profiles (94°C for 2 min, 11 cycles of 94°C for 20 sec, 56°C (declining 1°C each cycle) for 50 sec, 72°C for 30 sec, 25 cycles of 94°C for 2 min, 45°C for 50 sec, 72°C for 2 min, and a final extension of 72°C for 5 min).

### RNA extraction and sequencing

We homogenized individual aphids with a pestle in 500 μL of TRIzol (Invitrogen; Thermo Fisher Scientific, Inc., Waltham, MA, USA) and extracted total RNA using BCP (1-bromo-3-chloropropane; Life Technologies, Thermo Fisher Scientific, Inc., Waltham, MA, USA) with isopropanol precipitation. We used the Zymo RNA Clean & Concentrator kit (Zymo Genetics Inc., Seattle, WA, USA) to improve the purity and to remove gDNA using DNAse I. We then performed metatranscriptome Sequencing at Novogene (Novogene Corporation Inc., Sacramento, CA, USA). Library preparation was conducted using ribosomal RNA (rRNA) depletion by Illumina TruSeg Stranded Total RNA with Ribo-Zero Plus and NEBNext rRNA Depletion Kit (Zymo Genetics, Inc., Seattle, WA, USA). The libraries were sequenced to approximately 9 billion base pairs (bp) per sample with 150 bp paired-end reads on an Illumina NovaSeq platform. Raw reads were deposited into the NCBI Sequence Read Archive under BioProject ID PRJNA942253 with BioSample accessions SAMN33770905-SAMN33770908, and data accessions SRR23870213-SRR23870216.

### Microbial analysis using CZID

We assessed the success of ribosomal reduction in the metatranscriptome libraries using riboPicker [44] and the reference database SILVA_138 [45] (supplementary file reads_report.csv). We then used the CZ ID platform pipeline V7.1 (https://czid.org) [46], a cloud-based, open-source bioinformatics platform designed to detect microbes from metagenomic data. We removed host-specific reads (STAR host subtraction) using the *Acyrthosiphon pisum* genome [47], trimmed adapters using Trimmomatic [48], removed low-quality reads with PriceSeqFilter [49], and aligned the remaining reads to the NCBI NT and NR databases using Minimap2 [50] and Diamond [51]. In parallel, short reads were *de novo* assembled using SPADES [52] and mapped back to the resulting contigs using bowtie2 [53] to identify the contig to which each raw read belongs. We used the CZ ID water background model, which evaluates the significance (z-scores) of relative abundance estimates for microbial taxa in each sample. Potential bacterial reads were distinguished from contaminating environmental sequences by establishing z-score metrics ≥10, alignment length over 50 matching nucleotides (NT L ≥50), and a minimum of five reads per million aligning to the reference protein database (NR rPM ≥ 5). Potential viruses were established by z-score metrics of ≥1, NT L ≥50, and NR rPM ≥ 5 [46, 54, 55]. Bacterial and viral hits were confirmed with BLASTX and BLASTN manual searches. Only annotated non-host hits with revised Taxonomy IDs and BLAST-based match refinement were used for further analysis. The “Macrosiphum euphorbiae” project is viewable and searchable to anyone in CZ ID.

### Densoviral analysis using *de novo* assembly and Travis

We conducted an additional screening and viral genome assembly of potential Ambidensoviruses using *de novo* transcriptome assemblies obtained as follows. We used Trimmomatic v.0.39 [48] to trim the sequence adapters and filtered low-quality/complexity reads, and assessed for post-trimming quality using FastQC v.0.11.9 [56]. Then, we used Trinity v.2.14 [57] to *de novo* assemble the remaining reads. We used TRAVIS (v.20221029, https://github.com/kaefers/travis) to scan the assembled transcriptomes for Densovirus-like sequences. We built the reference database according to the currently accepted Densovirinae (ICTV, 29. Oct 2022, see supplementary file parvoviridae_reference_library.csv), extracted open reading frames between 100 and 2000 amino acids from the assembled transcriptomes, and screened using HMMER v3.3.1 [58], MMSeqs2 [59], BLASTP v2.12.0 [60], and Diamond v2.0.15 [51]. We set the e-value cutoff at 1 ×10^−6^, where applicable. All hits were again searched with Diamond against the non-redundant protein database (NCBI, downloaded on 29 Oct 2022).

### MeV-1 genome analysis

We used the CZ ID viral consensus genomes pipeline to build a consensus genome from the sample with MeV-1 present at high levels. In short, contigs were aligned to the reference MeV-1 genome (NCBI Entry KT309079.1) using minimap2 [50] and then trimmed using TrimGalore (Phred score <20) [61]. The consensus genome was generated with iVar consensus using a depth of five or more reads [62].

### MeV-1, MeV-2, and *Hamiltonella defensa* screening

Like all aphids, *M. euphorbiae* hosts an obligate heritable bacterial symbiont called *Buchnera aphidicola* that synthesizes amino acids missing from the aphid’s diet of plant phloem, and can also harbor several other facultative symbiotic bacteria (listed above) [43]. To screen for these microbes, we used 1 μg of total RNA extracted (as above) from each of the 23 adults collected during 2022 for cDNA synthesis with iScript cDNA synthesis kit (Bio-Rad Laboratories, Inc., Hercules, CA, USA). To screen for the Flavivirus Macrosiphum euphorbiae virus 1 (MeV-1), we used 100 ng of cDNA, the primers MevirF1 (5’-GTACACTTGCCTTACCTTACTGT-3’) and MevirR1b (5’-AACACGGGTCACGACCTTAG-3’), and the PCR conditions previously described [30]. To screen for the Ambidensovirus Macrosiphum euphorbiae virus 2 (MeV-2), we used 100 ng of cDNA, the MeV2-F (5’-CCGGATGACAAATCCCACGA-3’) and MeV2-R (5’-AATAGGCGCAGAGATGGACG-3’) primers, and the recommended PCR conditions [24]. In addition, we extracted DNA from colonized Me57 aphids (as above) and used 40 ng of genomic DNA to screen for MeV-2. The aphid Glyceraldehyde 3-phosphate dehydrogenase (G3PDH) was used as internal control (primers G3PDH_F (5’-CGGGAATTTCATTGAACGAC-3’) and G3PDH_R (5’-TCCACAACACGGTTGGAGTA-3’) [35]).

We used 200 ng of the cDNA previously synthesized for MeV-1 and MeV-2 screening and the protocols for *Hamiltonella defensa* PCR screening (as described above) to evaluate the proportion of field-collected aphids harboring this bacterial symbiont (supplementary file samples_metadata.csv). Furthermore, we used a non-parametric (Spearman) correlation to investigate the potential interaction between *Hamiltonella* and MeV-1.

### DNA extraction and sequencing

We pooled seven genetically identical adult unwinged aphids from cultivated lab line Me57 and isolated genomic DNA (gDNA) using a phenol/chloroform extraction. We then sheared the gDNA to approximately 20kb fragments using Covaris G-tubes (Covaris LLC., Woburn, MA, USA) at 4200 RMP for 1 minute, followed by tube inversion. For library preparation, we used the NEB Next PPFE repair kit with Ultra II end prep reaction (New England Biolabs, Ipswich, MA, USA) under recommended conditions and Nanopore ligation sequencing kit SQK-LSK110. For sequencing, we used a Nanopore R9.4.1 (FLO-MIN106D) flow cell and a MinION MIN-101B sequencing device (Oxford Nanopore Technologies, Oxford, UK). We ran the flow cell for 24 hours, followed by a wash with Flow Cell Wash Kit (EXP-WSH004); we then reloaded the flow cell with a second library prep and ran the sequencer for an additional 48 hours. We stopped the second sequencing run at 72 hours (∼22 Gbps of sequencing). In addition, we performed an additional 5.3 Gb of 150 bp paired-end sequencing to polish the assembly on an Illumina NovaSeq platform. DNA was extracted as above, and library prep and sequencing were performed by Novogene Inc. Raw reads were filtered for low quality and adapter contamination by Novogene Inc.

### *M. euphorbiae* whole genome assembly

We used Guppy (Oxford Nanopore Technologies) for base-calling and quality trimming raw reads. For the removal of *Buchnera* reads, we used minimap2 v.2.24 [50] in conjunction with SAMtools v.1.15.1 [63] to map our reads against the *Buchnera aphidicola* (strain *Macrosiphum euphorbiae*) genome (NCBI accession NZ_CP029205) and the corresponding plasmids (NCBI accession number NZ_CP029203 and NZ_CP029204). We only kept unmapped reads for aphid genome assembly. We assembled Nanopore reads using CANU v.2.0 [64] with an estimated genome size of 541 Mbp. We removed allelic variants from the assembly using the purge_haplotigs v.1.1.2 [65], first by mapping reads to the assembly using minimap2 v2.24-r1122 with Samtools v.1.15.1 and manually choosing cutoffs for haploid vs. diploid coverage based on a histogram plot (v -l 5 -m 27 -h 60), and then by purging duplicated contigs based on coverage level (-j 80 -s 50). For assembly polishing, we used the Illumina reads after quality assessment using FastQC V0.11.9 [56]. Then we used these reads to polish the purged assembly using Pilon v.1.24 with default parameters [66]. We used BlobTools2 [67] to identify remaining contaminating contigs. For this, we used blast results obtained from the BLASTN function against the NR database using blast plus v.2.12.0 [68], read coverage obtained by mapping the Illumina reads to the assembly using minimap2 v.2.24 [50], and GC content in this analysis. Based on these results, we removed all the short contigs with strong homology to the plant genus *Solanum* (which includes the tomato host plant species of *M. euphorbiae*) as we suspect these contigs were assembled from host plant contamination in the guts of sequenced aphids. We also removed two short contigs with homology to other bacterial contaminants such as *Escherichia coli* and *Pseudomonas* sp. Lastly, we removed a contig nearly identical to the pLeu plasmid found in *Buchnera aphidicola* and a small portion of two large contigs matching the *Buchnera* genome. The final annotation was assessed using BUSCO v.5.3.2 [69] with the MetaEuk gene predictor [70] implemented in galaxy.org, using the hemiptera_odb10 (2020-08-05) lineage dataset. The *M. euphorbiae* genome is available in NCBI with BioProject ID PRJNA942253 and BioSample SAMN33681650. The raw Nanopore (SRR23851809) and Illumina reads (SRR23919025) associated with the genome are available through the Sequence Read Archive, and the finished assembly is available with accession number JARHUA000000000.

### Characterizing endogenous viral elements in the *M. euphorbiae* genome

DNA Illumina raw reads were used as input to the CZ ID platform pipeline V7.1 (https://czid.org) and a z-score metrics of ≥1 and NT L ≥50 as described above [46, 54]. Additionally, to screen for actively transcribed Ambidensovirus-like EVES in the *M. euphorbiae* genome, we used BLASTN searches using the seven viral hits provided in individual Trinity contigs flagged by TRAVIS (supplementary file contigs_TRAVIS.fasta) against the genome scaffolds. All non-redundant hits from these searches with E-values < 1.10^−3^ were extracted and used in further analyses [33].

## Results

### Analysis of non-host sequences detected in single aphid

We used the pea aphid (*A. pisum*) genome to subtract host reads from our transcriptome data set. On average, 81.8% of the reads mapped to this host and were subtracted from further analysis (see supplementary file reads_report.csv). We then analyzed the remaining distribution of non-pea aphid reads, within a single *M. euphorbiae* aphid, as the overall proportion of reads assembled into contigs that could be assigned to bacterial, eukaryotic, and viral taxa (public project Macrosiphum euphorbiae at https://czid.org). Bacterial taxa dominated the microbial signature (Figure 1A), and as expected, the highest number of reads assembled into contigs matched the aphid obligate symbiont *Buchnera aphidicola* with over 45,000 reads per million aligning to the nucleotide database (NT rPM>45,000). Reads from an aphid facultative symbiont *Hamiltonella defensa*, were found in two samples (NT rPM>8,700). One sample (Me152) showed a strong signature of bacterial contaminants (*E. coli, Pseudomonas, Halomonas*, and *Agrobacterium*) commonly present in soil and plant surfaces.

**Figure 1.**
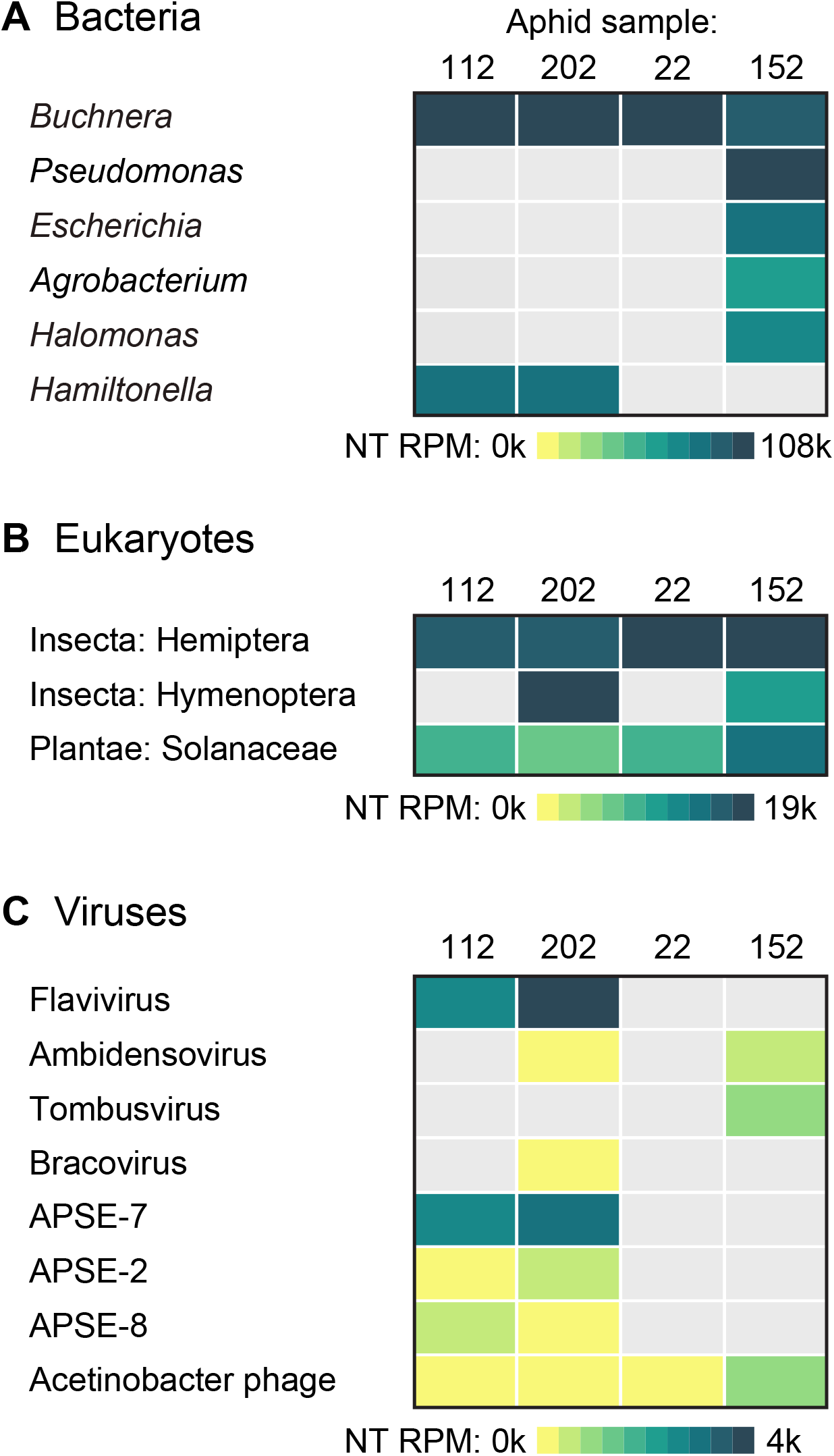
Details of the per aphid breakdown of non-host reads aligning to specific bacteria (**Figure 1a**), eukaryotic (**Figure 1b**), and viral (**Figure 1c**) taxa. Reads per million aligning to the nucleotide database (NT rPM) used as the quantitative metric in the heatmaps (see supplementary files heatmap_metrics.csv for metric details).

In terms of eukaryotes (Figure 1B), we found hits to Solanaceae, which includes the host plant species of *M. euphorbiae*, and Brachonidae parasitoid wasps (Insecta: Hymenoptera) in two samples (NT rPM>18,000). *M. euphorbiae* is known to be parasitized by hymenopterous wasps belonging to the superfamilies Ichneumonoidea (Braconidae) and Chalcidoidea [71]. In addition, there were some *M. euphorbiae* species-specific reads remaining, which did not map to the pea aphid reference genome but showed some homology to other aphid species (Insecta: Hemiptera).

We detected the presence of two insect-specific viruses in our metatranscriptome data (Figure 1C). The highest number of hits matched a previously described insect-specific Flavivirus, called Macrosiphum euphorbiae virus 1 (MeV-1), which we detected in two samples (NT rPM = 234 and 4055 for Me112 and Me202, respectively). We also detected viral hits to an insect-specific Ambidensovirus (Me202 and Me152; NT rPM>60). Other viral reads in our samples included a Bracovirus in one of the samples that was parasitized with the Brachonidae wasp (Me202; NT rPM=1) and a Tombusvirus (Me152; NT rPM=2.9), a family of plant pathogenic viruses with a single-stranded positive-sense RNA genome. Lastly, we detected two phage genera, the *Hamitonella-*specific phage APSE (NT rPM>310) in the same samples found positive for this symbiont (Me112 and Me202) and Acinetobacter phage (NT rPM 0.5-18), a bacteriophage largely prevalent in the environment [72].

### Comprehensive and quantitative analysis of insect-specific viruses

Using the CZ ID platform, we aligned five assembled contigs to the MeV-1 reference genome (NCBI accession KT309079) and found that they ranged between 85.8-97.2% nucleotide identity to the reference genome (Figure 2A). Our transcriptome retrieved 17,397 informative nucleotides allowing the assembly of a nearly complete genome for MeV-1. Our MeV-1 consensus genome has a coverage breadth of 79% and a coverage depth of 673.2x (NCBI accession OQ504571) (supplementary figure MeV1_coverage.tif). This single-stranded positive-sense RNA genome contains a single large ORF encoding a polyprotein of 7,333 amino acids, which is subsequently processed to generate structural and non-structural proteins [73]. Previous analysis indicated that the polyprotein motifs of MeV-1 helicase, methyltransferase, and RdRp are similar to domains in other *Flavivirus* (family Flaviviridae) [21, 30]. The characteristic secondary structures (RNA stem-loop) in *Flavivirus* genomes most likely contributed to the 5,283 missing bases in our MeV-1 consensus genome assembly [74].

**Figure 2.**
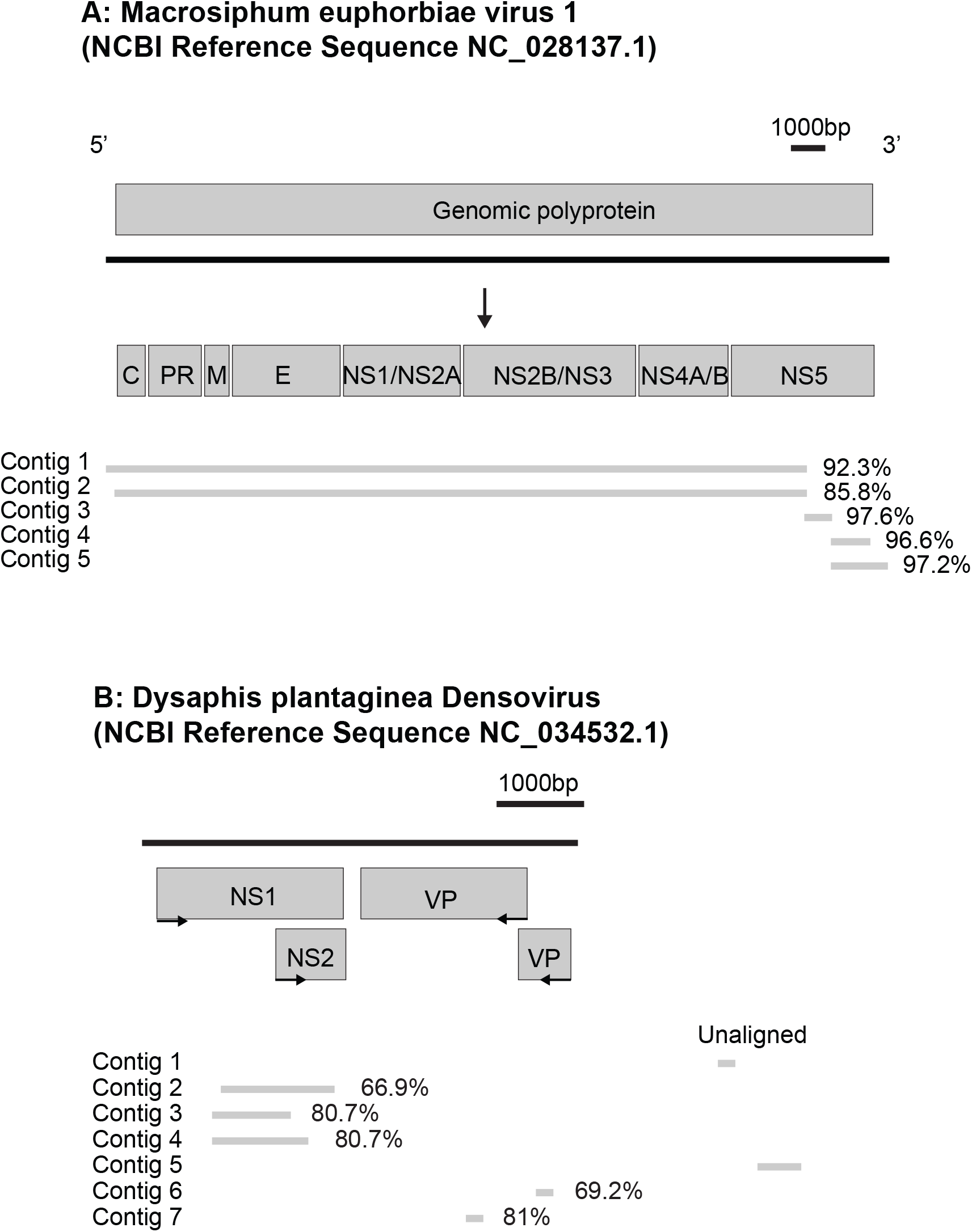
Assembled *M. euphorbiae* transcriptome contigs aligning to previously described insect Flavivirus (**Figure 2a**) and Ambidensovirus (**Figure 2b**) (see supplementary files contigs_CZID.fasta and contigs_TRAVIS.fasta for sequence details).

In addition, using the CZ ID platform, we detected two contigs with 80% similarity to the non-structural protein 1 (NS1) of Dysaphis plantaginea Densovirus (DplDNV), single-stranded DNA insect-specific *Ambidensovirus* (family Parvoviridae) (supplementary file contigs_CZID.fasta). Due to the lack of a publicly available genome or partial viral sequences of Macrosiphum euphorbiae virus 2 (MeV-2), an Ambidensovirus previously described in the same aphid species [24], we were not able to explore the homology between both viruses. Therefore, we conducted a more extensive analysis of our RNAseq data using TRAVIS, a consistency-based virus detection pipeline for sensitive mass screening of transcriptomic data directed toward Parvoviridae proteins. While degrees of sequence identity between Densovirinae (a subfamily of viral species exclusively infecting arthropods) is very low, viral species often express NS1 and VP proteins, which are useful for parvovirus phylogenetic inferences [75]. We used the seven viral hits provided in individual Trinity contigs flagged by TRAVIS (supplementary file contigs_TRAVIS.fasta) to identify the ORF orientation and similarity and to construct a hypothetical genome assembly using DplDNV as the closest reference available (Figure 2B). We found three contigs with 68.8% to 81.3% similarity to the non-structural ORF1 (encoding for the NS1 protein) and two contigs with 68.8% to 86.2% similarity to the structural ORF (encoding for the VP protein). None of the assembled contigs showed similarity to DplDNV ORF2 (encoding for the NS2 protein). We only detected 70% similarity with the ORF2 of a distantly related Ambidensovirus (NCBI accession AMG693112), which genomic organization differs from previously reported aphid densoviruses [21]. Importantly, all densoviral NS1-like sequences also showed a high nucleotide similarity (72-85%) to the pea aphid APNS-2 (NCBI accession NC_042493.1 and NC_042494.1), an EVE that contributes to wing phenotypic plasticity in this species [35].

### Insect-specific virus frequency in natural populations

To further investigate the infection frequency of MeV-1 and MeV-2 infections in natural populations, we used a PCR approach to screen 23 individual adult aphids collected during 2022 as well as aphids from our colonized *Macrosiphum* line (Me57). We found only 13 field-collected aphids positive for MeV-1 (54.2%) and 21 aphids (87.5%) positive for MeV-2, including the colonized individuals (Figure 3). We also tested the cDNA of field-collected aphids (previously screened for MeV-1) for the presence of *Hamiltonella defensa* and found that 54.2% of the aphids (n=13) were harboring this bacterial symbiont. We found that 41.7% of individuals (n=10) shared a co-infection between this Flavivirus and *Hamiltonella* (Figure 3), but this association was not statistically significant (p-value= 0.078; r= 0.375).

**Figure 3.**
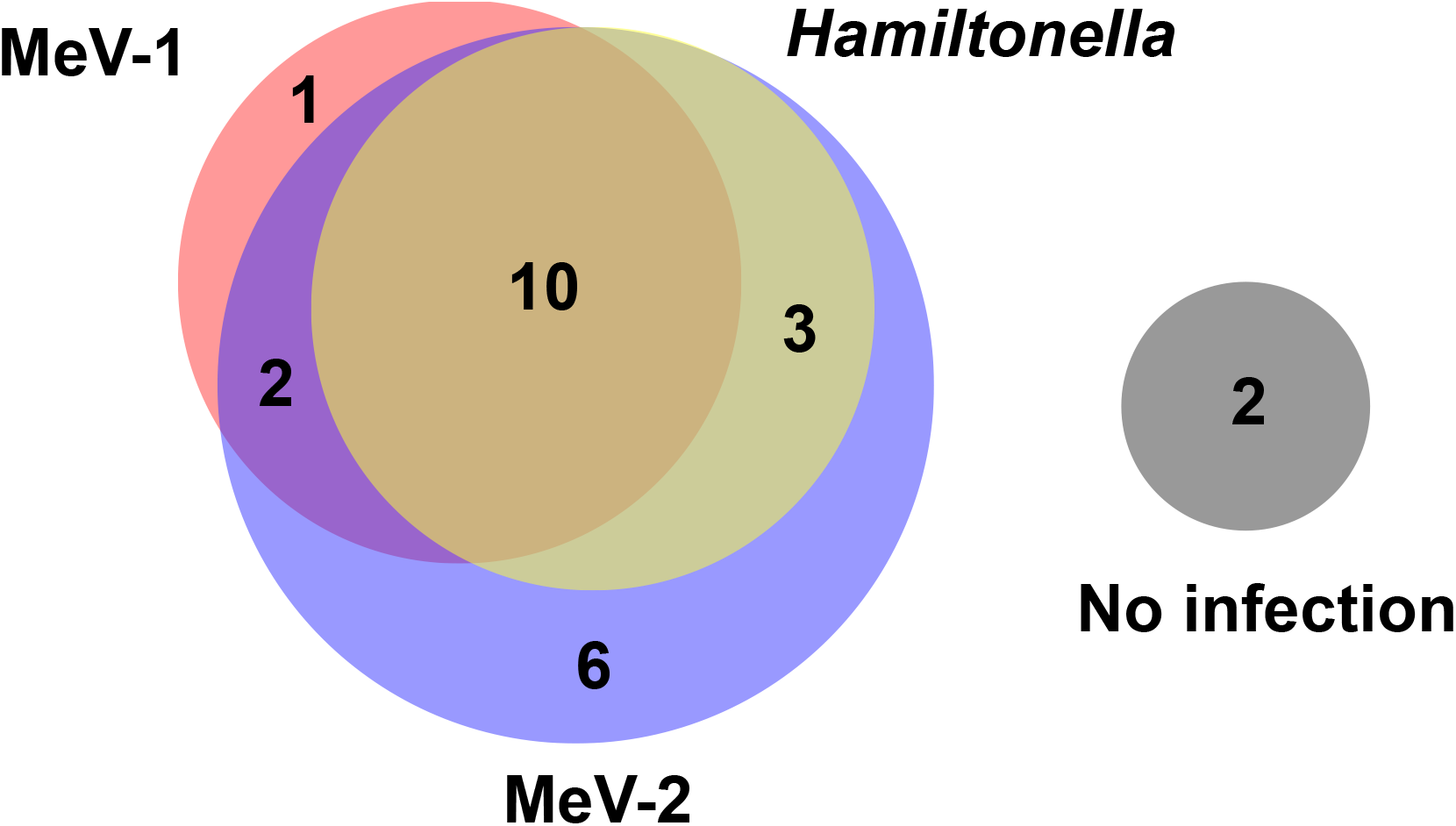
Frequency of Macrosiphum euphorbiae virus 1 (MeV-1), Macrosiphum euphorbiae virus 2 (MeV-2), and *Hamiltonella denfesa* infections in wild-collected (n=23) and colonized (n=1) aphids. All samples tested using cDNA for PCR screenings.

### Genome sequencing for analysis of endogenous viral elements (EVEs)

Our laboratory line (Me57) was found to be PCR positive for MeV-2, and we then used DNA sequences from a pooled sample of Me57 aphids to look for viral reads. We used the CZ ID platform as above to identify viral taxa using the Illumina DNA reads from our colonized Me57 aphid line. Surprisingly, we detected only a single contig with a low number of Ambidensoviral hits (NT rPM>0.329), which also showed 79.0% similarity to the DplDNV NS1 viral protein and 84.34% similarity to an uncharacterized genomic transcript in pea aphids (NCBI accession XM_029492170.1). Since both of our transcriptome and genomic data were unable to recover a complete or near-to-complete Ambidensovirus genome, we then suspected that these viral reads could correspond instead to actively transcribed EVEs, as previously reported in other closely related aphid species [33, 35].

To determine with certainty whether the Ambidensovirus hits found in our transcriptome data corresponded to an actively transcribed EVE, we assembled the first *M. euphorbiae* genome publicly available. We obtained a total of 4,223,264 nanopore reads (at an average of 5.21kb) and 35,578,886 Illumina reads (PE 150bp) from sequencing. After assembly, haplotig purging, polishing, and manual removal of plant and bacterial contigs, our assembly contained 2,176 contigs with an N50 length of 665kb and a total length of 545.7 Mb (Figure 4A). *M. euphorbiae* has a similar GC content (29.96%; Figure 4B) to other sequenced aphids (e.g., *A. pisum* at 29.6%, *M. persicae* at 30.1%, and *A. glycines* at 27.8%) [76, 77]. The size of our assembly is close to a recent estimation of the *M. euphorbia* genome size based on flow cytometry which was estimated at 531.7 Mb [76]. Similarly, an analysis of single-copy orthologs showed our assembly contains 98.5% complete BUSCOs, with 94% present in single copies and 4.5% duplicated (Figure 4C). An additional 1.2% of BUSCOs are fragmented, and 0.3% are missing. Together these results suggest that this draft of the genome is highly complete.

**Figure 4.**
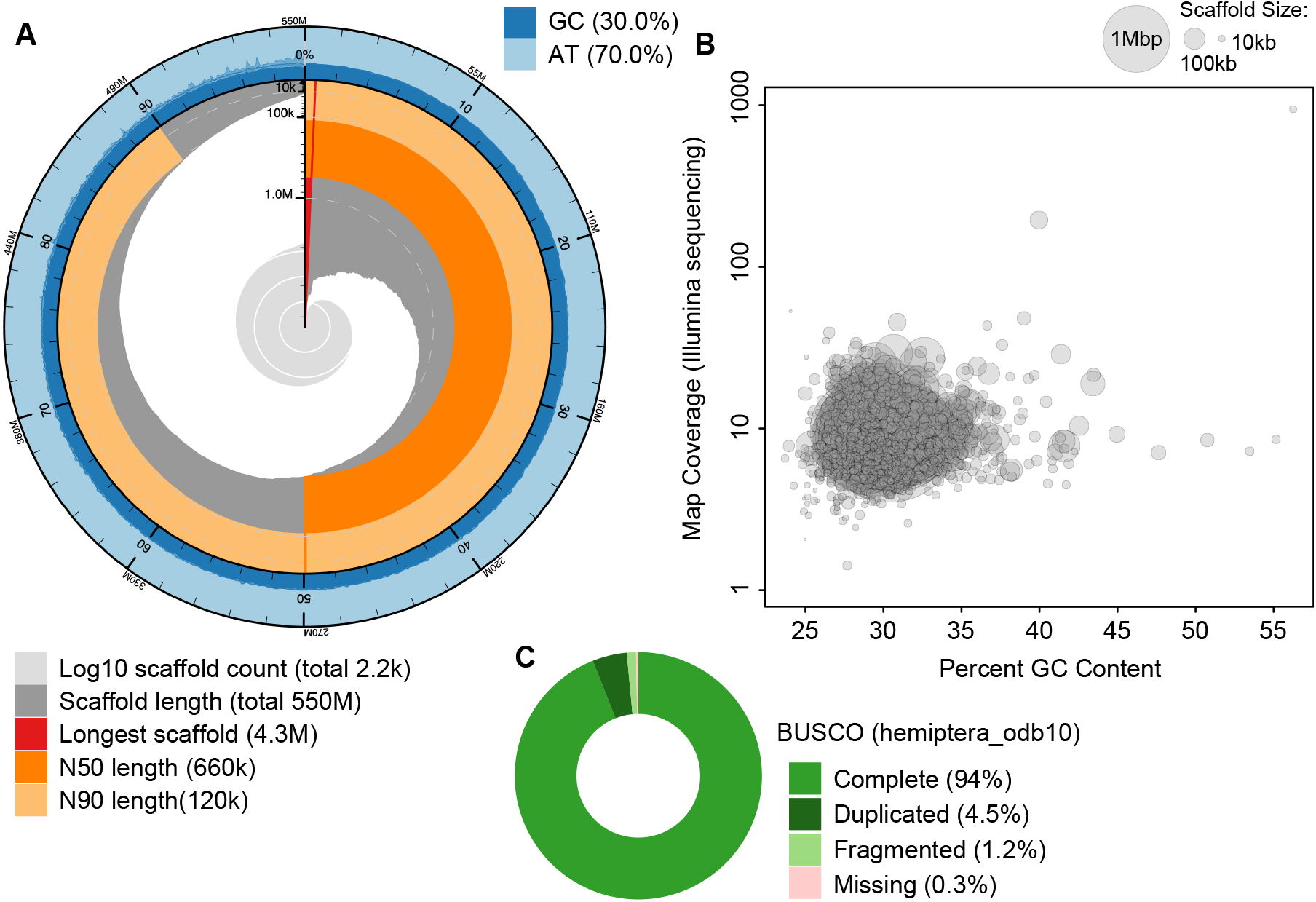
M. *euphorbiae* genome assembly metrics (**Figure 4a**), GC content and coverage (**Figure 4b**), and BUSCO metrics (**Figure 4c**).

We used the genome as a reference to screen for the seven individual Trinity contigs flagged by TRAVIS as potential Ambidensovirus in our previous analysis. Initially, we selected hits with E-values < 1.10^−3^ [33]; however, most of the 3,044 hits represent shorter sequences rather than the actual transcript length (see supplementary file expressed_densoviral_EVEs.csv); therefore, we restricted the search to matches consistently to the entire length of each transcript and E-values=0 (Table 1). No full-length hits in the genome were found for the two largest viral contigs assembled from transcriptome data (contig3 and contig4); instead, the best hits for these two contigs corresponded to 16-17% of the total length. In insects, the EVE repertoire varies between distinct populations of a given species and, in some cases, even between individuals within the same population [78]. This phenomenon explains why all the field aphid samples (n=3) that tested negative for MeV-2 by PCR amplified a product of approximately 500 bp, which is about half of the expected size reported for the primers used. Given that the genome assemblies and RNAseq data sets were derived from different aphid strains, it is not surprising the wide range of partial-length Ambidensovirus hits obtained in our analysis. However, we are confident that five full-length viral transcripts are constitutively expressed from three regions of the *M. euphorbiae* genome (tig00030708_pilon, tig00029914_pilon, and tig00027226_pilon).

**Table 1.** List of Ambidensoviral transcripts and the corresponding integrations in *M. euphorbiae* genome.

| Transcriptome contig | Transcript length | Percentage of identical sites | Hit end | Hit start | Genome contig | Query end | Query start |
| --- | --- | --- | --- | --- | --- | --- | --- |
| Travis_contig1 | 783 | 96.70% | 783 | 1 | tig00030708_pilon | 198038 | 197258 |
| Travis_contig1 | 783 | 98.90% | 1 | 783 | tig00029914_pilon | 60345 | 59559 |
| Travis_contig2 | 466 | 96.20% | 416 | 1 | tig00030708_pilon | 198433 | 198018 |
| Travis_contig2 | 466 | 99.80% | 1 | 466 | tig00029914_pilon | 59579 | 59114 |
| Travis_contig3 | 2155 | - | - | - | - | - | - |
| Travis_contig4 | 2878 | - | - | - | - | - | - |
| Travis_contig5 | 1174 | 99.90% | 1 | 1174 | tig00030708_pilon | 92758 | 91585 |
| Travis_contig6 | 1040 | 99.80% | 1 | 1040 | tig00030708_pilon | 85562 | 84525 |
| Travis_contig7 | 635 | 100.00% | 635 | 1 | tig00030708_pilon | 86191 | 85557 |
| Travis_contig7 | 635 | 84.90% | 635 | 1 | tig00027226_pilon | 138266 | 137632 |

## Discussion

RNAseq is becoming an essential tool for virus discovery. Our study illustrates how endogenous viral elements in insect genomes can be an obstacle to using RNAseq for characterizing viral diversity in arthropods. We used rRNA-depleted RNAseq along with bioinformatic tools to characterize the virome of an important insect pest species, the potato aphid *Macrosiphum euphorbiae*. Our analysis found two insect-specific viruses from the families Flavivirus and Ambidensovirus described in previous RNAseq studies. However, by sequencing and assembling the genome of this insect using long-read sequencing, we found that the Ambidensovirus is a transcriptionally active EVE rather than an exogenous virus. Endogenous viral elements encoded in the host genome are abundant in arthropod genomes, and thus EVE sequences in RNAseq studies are an important consideration for future studies of viral diversity in arthropods.

Densoviral EVEs have been shown to be transcriptionally active in two other aphid species: *Myzus persicae* and *A. pisum*. In pea aphids, two copies of a transcribed densoviral non-structural protein (termed the “APNS” genes) were found to be upregulated in response to crowded conditions and to be functionally linked to the plastic production of wings [35]. These genes had close homology with the non-structural genes of Dysaphis plantaginea densovirus (DplDNV), which, when infecting rosy apple aphids, causes them to be winged [79], suggesting the function of these viral genes had been conserved after endogenization. The transcribed EVEs we found in *M. euphorbiae* have significant homology to the pea aphid APNS genes, and it seems likely that these genes may also be playing a role in wing plasticity in *M. euphorbiae* though additional data is needed.

Our study contributes to the growing list of sequenced aphid genomes, which together show that transcribed densoviral EVEs are common in this insect group [33, 35, 37, 80]. Most identified EVEs in insect genomes correspond to unclassified single-stranded RNA viruses and viruses belonging to the families Rhabdoviridae and Parvoviridae [78]. Unlike RNA viruses, which may produce abundant short mRNAs that favor virus endogenization [20], Parvoviruses undergo a double-stranded DNA intermediate during nuclear replication, which along with the endonuclease activity of NS1 and the eukaryote double-stranded break repair mechanism may largely favor endogenization of this virus family [81, 82]. Previous studies have estimated that around 10% of the parvoviruses described in animals are likely integrated into host genomes, but in most cases, the EVE status remains uncertain due to unavailable or incomplete genomes for those species in which transcriptome data is available [75]. Multiple recent studies have described the presence of “new” densoviruses in aphid’s transcriptome [23, 26, 83]; however, our combined transcriptomic and genomic analyses suggest that some of those viral transcripts may likely correspond to actively transcribed EVEs instead of heritable exogenous viruses infecting aphids at very high rates.

Last, our study shed light on the biology of MeV-1, an insect-specific Flavivirus (family Flaviviridae), previously characterized by RNAseq studies of *M. euphorbiae* populations collected in France [30]. We found that this virus, contrary to previous reports, is present in a North American population of *M. euphorbiae*, and we found that it is highly prevalent. By assembling the genome of MeV-1 from our RNAseq data, we found that our local population is infected with a potentially distinct viral strain from previous studies. No obvious infection symptoms or abnormal phenotypes were observed in MeV-1-infected *M. euphorbiae* adults, and future studies are needed to determine what phenotypic effects this virus has on its host. Other heritable viruses have been found to interact with the secondary symbiotic bacteria found in aphids [84-86] but we did not find significant patterns of co-infections with the bacterial symbiont *Hamiltonella defensa*.

EVEs are common in insect genomes, and our results highlight this widespread challenge in studying insect viromes. Our study further emphasizes how combining sequencing methodologies is necessary to overcome the potential pitfalls of only RNAseq-based viral discovery. Careful consideration of the biological characteristics and genome structure of viruses discovered through RNAseq is essential [87]. In aphids and other widely study systems, the development of cultured cell lines is also imperative to isolate viral species described by sequence-based methods, to characterize viral replication, and for use in large-scale virus production that will facilitate future investigation of the complex interaction of aphid viruses and their hosts [21].

The relatively high transcription level of some EVEs suggests that viral integration may have important biological implications for the fitness of aphids. Likewise, uncovering the phenotypic effects of accurately described insect-specific viruses may also show promising targets for alternative control strategies of agriculturally destructive organisms while providing important foundational resources in the study of host-virus dynamics. Research efforts need to be done on the evolutionary dynamics of heritable viruses to better understand how they are acting as hidden drivers of host phenotypes.

## Supporting information

supplementary figure MeV1_coverage.tif

supplementary file expressed_densoviral_EVEs.csv

supplementary file parvoviridae_reference_library.csv

supplementary file reads_report.csv

supplementary files heatmap_metrics.csv

supplementary file samples_metadata.csv

supplementary file contigs_TRAVIS.fasta

supplementary files contigs_CZID.fasta

## Data accessibility

All data is available through CZ ID (Macrosiphum euphorbiae project), NCBI BioProject ID PRJNA942253, or included as a supplemental file.

## Authors’ contributions

PRL and BJP conceived of the project. PRL and BJP wrote the manuscript.

PRL, WB, SK, BJP carried out the data curation, bioinformatic analysis and software validation. PRL, WB, MMM carried out the molecular work. BJP contributed to funding acquisition. All authors edited the manuscript and approved the final version for submission.

## Acknowledgments

Thanks to Keertana Tallapragada, Georgina Aitolo, and Mariam Sirag for help with insect stock maintenance. The University of Tennessee Knoxville’s Office of Innovative Technologies provided access to the ISAAC-NG computational cluster.

