## Supplementary figures and images for "Untangling an insect virome from its endogenous viral elements"

### supplementary figure MeV1_coverage.tif

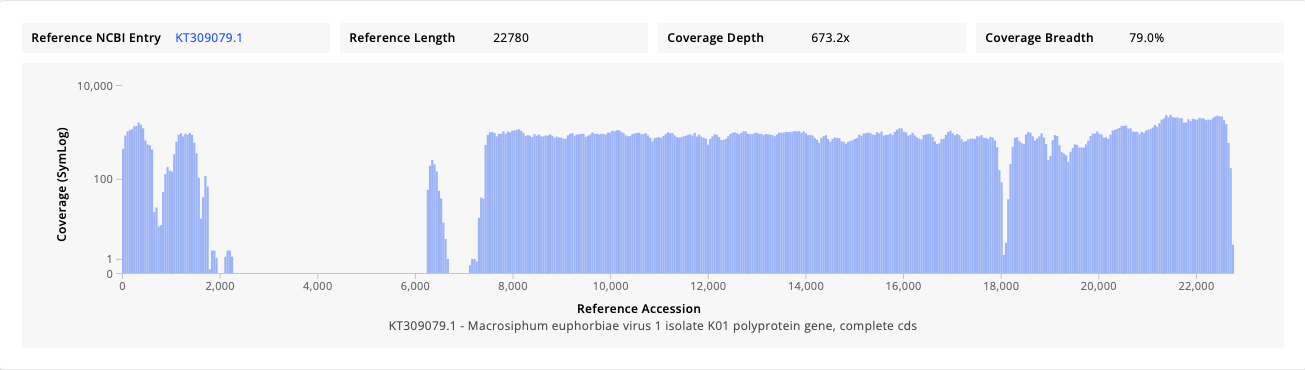
